# Population extinctions driven by climate change, population size, and time since observation may make rare species databases inaccurate

**DOI:** 10.1101/506006

**Authors:** Thomas N. Kaye, Matt A. Bahm, Andrea S. Thorpe, Erin C. Gray, Ian Pfingsten, Chelsea Waddell

## Abstract

Loss of biological diversity through population extinctions is a global phenomenon that threatens many ecosystems. Managers often rely on databases of rare species locations to plan land use actions and conserve at-risk taxa, so it is crucial that the information they contain is accurate and dependable. However, climate change, small population sizes, and long gaps between surveys may be leading to undetected extinctions of many populations. We used repeated survey records for a rare but widespread orchid, *Cypripedium fasciculatum* (clustered lady’s slipper), to model population extinction risk based on elevation, population size, and time between observations. Population size was negatively associated with extinction, while elevation and time between observations interacted such that low elevation populations were most vulnerable to extinction, but only over larger time spans. We interpret population losses at low elevations as a potential signal of climate change impacts. We used this model to estimate the probability of persistence of populations across California and Oregon, and found that 31%-56% of the 2415 populations reported in databases from this region are likely extinct. Managers should be aware that the number of populations of rare species in their databases is potentially an overestimate, and consider resurveying these populations to document their presence and condition, with priority given to older reports of small populations, especially those at low elevations or in other areas with high climate vulnerability.

## Introduction

Population extinctions are a major threat to plants, leading to range contractions, fragmentation and isolation [e.g., 1-4], which together reduce the abundance of species. As Darwin [5] pointed out, rarity is a precursor of extinction. Orchids in particular face a global conservation risk with high species diversity but also a high rate of species that are rare or threatened with extinction [6-12], and rare orchids are likely to need aggressive conservation actions to prevent their extinction [13]. Nearly half of the genus *Cypripedium* may be threatened and in need of protection if the species are to survive in the wild [14]. Therefore, accurate assessments of the number of populations of a rare species and its major threats are crucial to conservation planning and resource allocation for recovery actions [15, 16].

Several processes can contribute to rare plant population extinctions, including habitat loss, interactions with invasive species, changes in disturbance frequency, etc. [17]. Climate change in particular is affecting species ranges globally [18], with organisms shifting toward higher latitudes [19] and elevations [20]. For example, plant ranges in western Europe have moved upslope at 29 m/decade over the last century [21] and in California at similar rates [22]. Climate change effects on temperature and moisture may threaten plant diversity in Europe, especially in mountains [23]. Low-elevation populations of organisms can be especially at risk of extirpation as climatic conditions change and force upslope range shifts [24]. Any contraction in the range of a rare species can have significant effects on its long term conservation and viability.

The number of individuals present can also affect the viability of plant populations, with small populations having greater risk of extirpation. In general, the extinction probability of a population increases as population size decreases [25, 26]. Small populations may be at greater risk of extinction because of several factors, including losses in reproductive individuals [27], Allee effects [28], declines in seed production [29] and viability [30], loss of genetic diversity [31] and accumulation of genetic load [32], and demographic stochasticity [33]. In empirical studies that surveyed the same locations of multiple plant species over several years in Germany [4] and the Swiss Jura Mountains [34], extinction rates were found to be higher for small populations. And although population size may be a strong predictor of population vulnerability, passage of time can compound the likelihood of extinction because as more time passes in stochastic environments the chances that a population will fall to zero increase [25, 26].

Taken together, climate change, population size, and time since observation create considerable uncertainty regarding the current status of wild plant populations recorded in various rare species databases. Several US agencies and organizations (e.g., US Bureau of Land Management, US Fish and Wildlife Service, US Forest Service, NatureServe, state Natural Heritage Programs) maintain databases of rare plant occurrences and many of these occurrences may not have been visited recently. Therefore, the number of populations in the wild of some species could be smaller than the number listed in databases due to extinctions that have not yet been detected. Increasing our ability to estimate the number of populations that remain extant or have gone extinct in these data bases will improve conservation planning for rare species. We used information on repeated surveys in California and Oregon for a rare but widespread orchid, *Cypripedium fasciculatum* (clustered lady’s slipper), to test the hypothesis that extinction probability is affected by elevation, population size, and time since observation. We applied the resulting model to populations in Oregon and California in the Geographic Biotic Observations (GeoBOB) data base maintained by the US Bureau of Land Management and the US Forest Service Natural Resource Information System (NRIS-Terra) to estimate the number of populations that are still extant.

## Materials and methods

### Study species

*Cypripedium fasciculatum* (clustered ladies slipper; Figure 1) occurs in scattered population centers in western North America in California, Oregon, Washington, Idaho, Montana, Utah, Wyoming and Colorado. In California and Oregon, this taxon occurs predominantly in the Klamath-Siskiyou Mountains and Sierra Nevada Mountains. The United States Forest Service (USFS) considers it to be a Sensitive Species and the Bureau of Land Management (BLM) lists it as a Bureau Sensitive Species, and it is considered globally secure because of its widespread geographic range and abundance in some states [35]. In California and Oregon the species is most often found on north facing slopes in mixed coniferous forests of >60% canopy closure [36]. *Pseudotsuga menziesii* is the most common associated tree, but other frequently noted forest components include *Abies concolor, Cornus nuttallii, Pinus lambertiana*, and *Calocedrus decurrens*. Clustered lady’s slipper is known to occur in California and Oregon at elevations from about 180 to nearly 2000 m. The species has a complex life-history and depends on specific mycorrhizal fungi [37], which may affect its seed germination and growth. Mycorrhizal fungi may determine where and in which specific habitats this orchid can grow and how it responds to disturbance, but little information is available on the fungi, their requirements, associated tree species, and their function in forest ecosystems [36].

**Figure 1.**
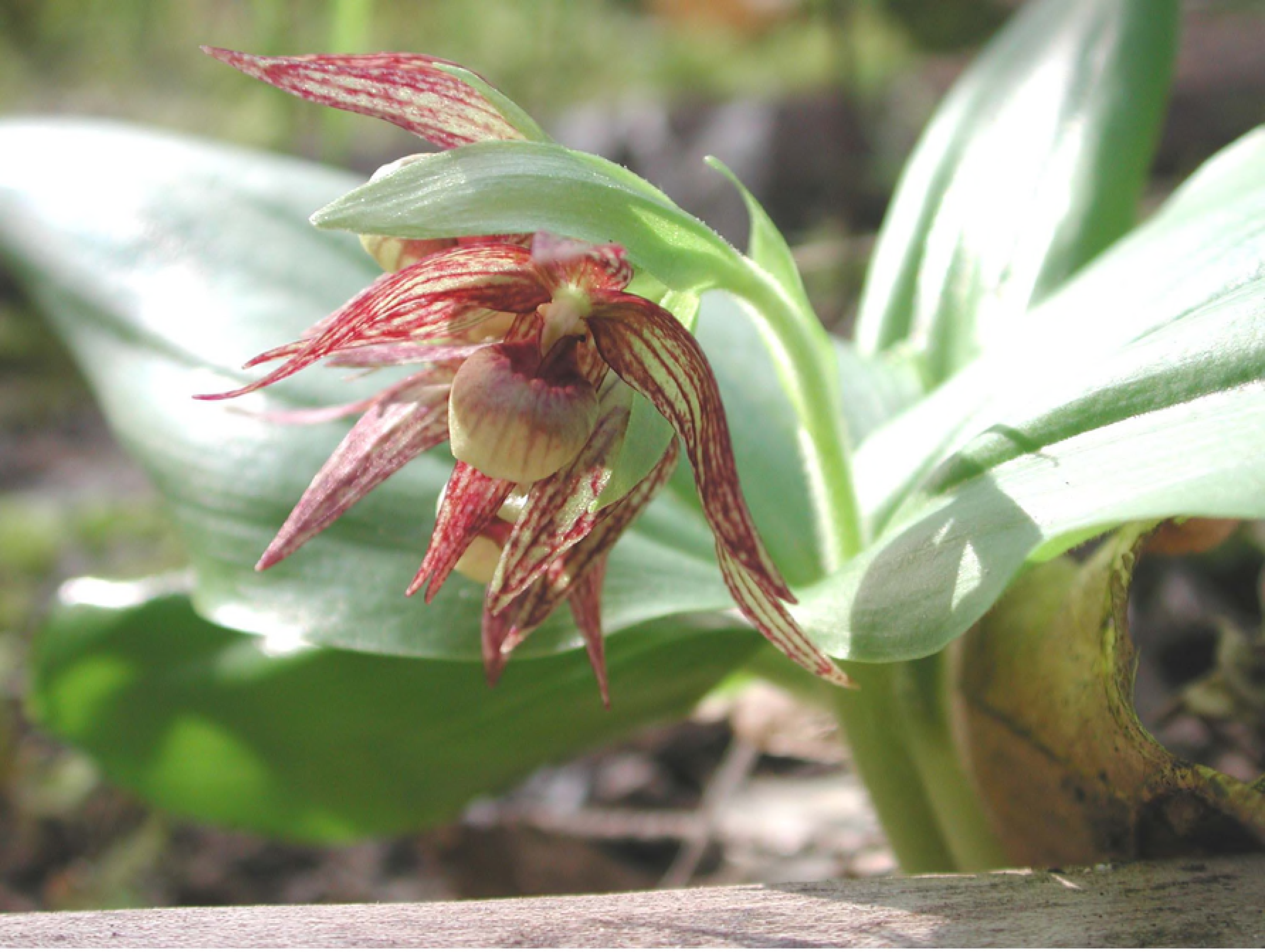
*Cypripedium fasciculatum* (clustered lady’s slipper).

### Data sources

We compiled repeated-survey data from multiple sources to test for effects of elevation, time between surveys, and population size on extinction probability. The sources of these resurvey data were from an assessment of the conservation status of *C. fasciculatum* in California that reviewed available records (78 sites) for the species throughout that state [36] and from repeated surveys in southwestern Oregon (158 sites) conducted on federal lands. Both resurvey data sources (236 populations combined) included sites revisited at last once and documented site location, elevation, population size, and time between surveys. We used information on population size from the first survey, and time between surveys was calculated as the number of years between the first and last (most recent) survey. The last survey was used to score each population as either extant or extinct (no individual plants found at the site). The time between surveys ranged from 1 to 29 years. While most observers censused populations, some estimated population size, and when this occurred we used the highest integer reported for a population during the first survey. For example, if 50-100 plants were reported, we used 100. If the number was vague (e.g., 75+, >30, or ca. 50) we used the actual integer listed (75, 30, or 50, respectively). Populations used in the analysis varied in size from 1 to 1084 individuals. *C. fasciculatum* plants that were single stems or clumps were considered individuals [following 40].

### Population Viability Analysis

We used a generalized linear model with quasibinomial errors to estimate extinction probability. The response variable was population status at the most recent visit (a binomial response, either extinct or extant) and independent variables were size of the population at the first survey, elevation of the population, and number of years between the first and last survey. All analyses were performed in R 3.3.2 [R Core Development Team, www.cran-r.org].

### Estimating extant population number

To estimate the number of populations of *C. fasciculatum* recorded as still extant in the GeoBOB and NRIS-Terra databases for California and Oregon, we applied our statistical model for predicting extinction probability to each of the 2896 populations recorded based on their elevation, size and years since the last survey. To estimate uncertainty, we bootstrapped the parameters in the model from our resurvey data set of 236 populations by randomly selecting 236 populations from this group, with replacement, and estimating the generalized linear model parameters. For each bootstrapped set of parameters, we calculated the extinction probability of each population in the GeoBOB and NRIS-Terra databases, summed those probabilities to estimate the number of extant populations, and repeated this bootstrap process 10,000 times to estimate 95% confidence limits. We performed this analysis in R 3.3.2.

## Results

### Population Viability Analysis

Of the 236 populations in our data sets, we found 34% were no longer present when revisited. Elevation, time between surveys, and population size were each significant factors for predicting extinction probability of populations (Table 1). Probability of extinction was best explained by all of these factors, including a significant (p=<0.001) interaction between elevation and years between surveys. The general linear model suggested that populations at lower elevations were more likely to go extinct than high elevation populations, but only as the length of time between surveys increased (Figure 2, right). Small populations had a greater probability of extinction than large populations, and extinction probability was near zero for populations with >100 individuals (Figure 2, left), regardless of the length of time between samples. Further, extinction probability increased as the time between surveys increased, most notably for smaller populations.

**Table 1.** Generalized linear model for factors affecting the probability of population survival for *C. fasciculatum*.

| Factor | Estimate | Standard Error | t value | P |
| --- | --- | --- | --- | --- |
| (Intercept) | 1.75 | 1.03 | 1.70 | 0.091 |
| Starting population size | 0.09 | 0.02 | 4.36 | <0.001 |
| Years between surveys | -0.38 | 0.1 | -3.81 | <0.001 |
| Elevation | -0.0002 | 0.0003 | -0.713 | 0.476 |
| Years between surveys<br>X Elevation | 0.00008 | 0.00003 | 2.76 | 0.006 |

**Figure 2.**
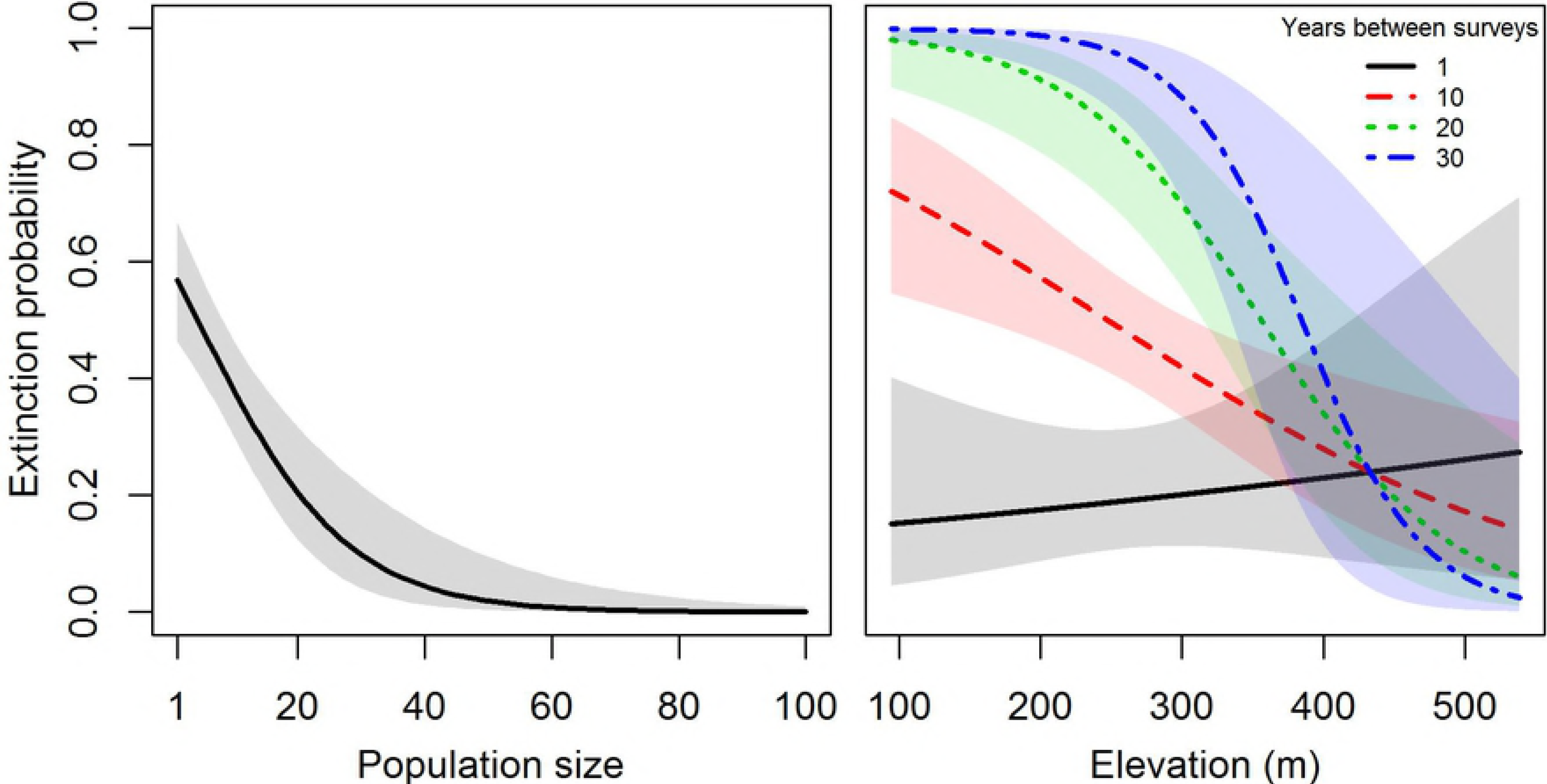
Extinction probability as a function of population size (left) and the interaction between elevation (m) and time (years) between surveys (right, with each line representing an example of a specific time interval between surveys. Shadings around each line represent 95% confidence intervals.

### Estimating extant populations

A total of 2415 populations with one or more plants were reported in the GeoBOB and NRIS-Terra databases for Oregon and California. An additional 426 populations were reported as already extinct by 2016. Populations in that database ranged in size from 1 to 1859 individuals, with a mean population size of 27 (95% CI ± 1.6). We estimated that of the 2415 populations reported as extant, only 1,349 (95% bootstrapped quantiles: 1213-1486) were likely still present. This is equivalent to an overall extinction rate of 44% (95% bootstrapped quantiles: 38%-50%). The predicted probability of population survival varied widely across the landscape in California and Oregon, with some population centers showing greater potential for population extinction than others (Figure 3). For example, populations in southwestern Oregon had a predicted extinction rate of 60% (53% – 67%) of 1258 reports compared to 27% (19% - 35%) of 1157 records in California. This difference was driven in our model by the generally lower population sizes in Oregon (mean: 12.7 95% CI: ± 1.5) than California (37.9 ± 6.6) and lower elevations of populations in Oregon (757.2m ± 13.4m) than California (1319.2m ± 15.4m). Years between observations did not differ between states, averaging 15.4 years overall (± 0.39).

**Figure 3.**
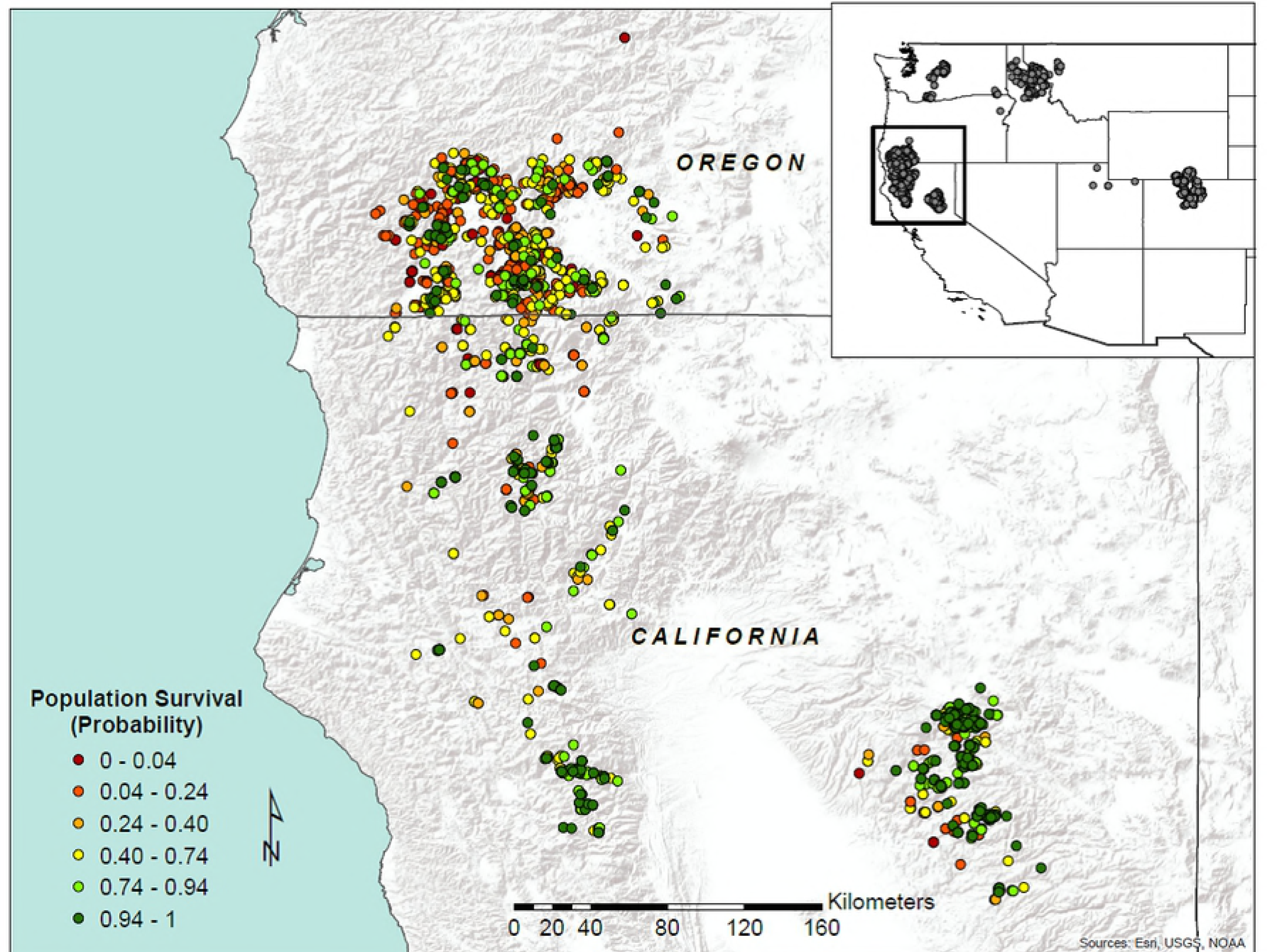
Distribution of *Cypripedium fasciculatum* in the western United States (inset), with map of California and Oregon showing the probability of persistence estimated from elevation, population size, and time since observation.

## Discussion

We found that elevation, population size, and time between surveys predicted extinction in *Cypripedium fasciculatum*. When these factors were used to model the persistence of wild populations, we found that only 56% of populations reported in the GeoBOB and NRIS-Terra databases for California and Oregon are likely still present on the landscape. Extinction rates are predicted to be higher in Oregon than in California, primarily due to the lower average population size and elevation there. Our results suggest that negative impacts from climate change might already be apparent for *C. fasciculatum* through extinction of low elevation populations. Loss of low elevation populations may be expected when climates warm to the point that populations can no longer survive in the hotter portions of their range. For example, loss of butterfly species at low elevations has been attributed to warming trends in Spain [39]. Our findings with *C. fasciculatum* are generally consistent with orchid responses to climate change in North America and elsewhere. Documented declines of species in the Orchidaceae in eastern North America appear to be related, at least in part, to an inability of these species to alter their phenology, particularly flowering time, as climate has warmed over the last century and a half [40]. Climate change appears to be a threat to orchids in Mexico [41], and orchids in general appear to be highly vulnerable to climate change in China [42]. In contrast, orchids were more likely to increase abundance in Mediterranean France from 1886-2001 compared to many other plant taxonomic groups [43]. Precipitation appears to be a strong driver of plant survival in *C. reginae* [44], making the species vulnerable to changes in regional climate. And it is clear that climate has changed recently and is forecasted to change further in California and Oregon, in part due to warming and drying that, when combined, exacerbate moisture deficits and increased evaporative demand [e.g., 45].

Many orchid species have populations with a wide range of sizes [46], and small average population sizes are common. In the GeoBOB and NRIS databases of 2415 populations of *Cypripedium fasciculatum* in California and Oregon that we reviewed, the average population size was 25 individuals. The average population size of *C. kentuckiense* is 40 individuals, *C. calceolus* in Europe generally has populations with fewer than 100 plants, and *C. dickensonianum* occurs as small colonies or individuals [47]. As population size declines in orchid species, gene flow by pollen may decline [48], inbreeding may increase [49], pollination, fruit set and seedling recruitment may decrease [50], genetic drift may increase [51], and genetic diversity may decline [52]. Transition matrix models of *C. calceolus* [53] indicate extinction probability over a 100 year period in populations with 10 plants is 37%, and in populations with 5 plants it increases to 67% without disturbance. In populations where flowers are removed or plants are dug up, extinction probability rapidly approaches 100%. The typically low population size in *C. fasciculatum* was a major contributor to the high rate of predicted extinctions we have shown for the species.

Population extinction probability was associated with time between surveys in *C. fasciculatum*. In stochastic environments, even populations with stable intrinsic population growth rates are vulnerable to extinction, and this vulnerability increases with time [25, 26]. In populations with declining growth rates, the rate of extinction will be even faster. Therefore, as time between surveys increases, population extinction should also increase, especially for small populations. Surprisingly, time between surveys had no significant effect on probability of extinction in eight rare plants in Germany [4], but the study was conducted over a relatively short period (ten years).

Resurveys of plant populations and communities can provide substantial insights into the nature and causes of changes that occur in the natural world over time [54-57]. Even so, there are some limitations to our estimates of extinction probability of *C. fasciculatum* in this study. Repeated surveys may fail to relocate previously documented populations even when they are still present [58-60] if the survey is not sufficiently thorough. The datasets we used contained information on population resurveys that were carefully conducted with precise location information, but the possibility remains that some extant populations may have been missed. This could be aggravated by individual plant dormancy, which would make plants very difficult to detect during surveys, and if all plants in a population were dormant at the same time – a possibility that increases as population size declines – whole extant but dormant populations could be falsely classified as extinct. Dormancy above ground is not uncommon in terrestrial orchids [61], including *Cypripedium* [38, 62-66]. *Cypripedium reginae,* for example, may be dormant for up to four years [44]. On the other hand, dormancy is associated with decreased orchid reproduction [67] and survival [68], and if all individuals in a population were dormant, the population might already be close to extinction. These factors suggest that although we could have overestimated extinction probability [58] due to dormancy, this same dormancy could suggest increased plant vulnerability. Either way, we are unable to quantify this potential bias in our results given the available data.

Because orchids depend on fungi, at least in the early stages of plant development, the presence of appropriate fungi and the environmental factors that affect them may in turn determine the growth and survival of many orchids [69], including *C. fasciculatum* populations. Soil and topography, and especially temperature and moisture are the most important factors that control orchid distribution and survival [70], and this may be due to the influence of these factors on mycorrhizal fungi. *Cypripedium* spp. are associated with fungi in the Sebacinaceae, Ceratobasidiaceae, and especially the Tulasnellaceae [37]. The degree of specificity of orchids with fungi is significant because orchids with highly specific associations may be more sensitive to disturbance and environmental change than generalist species [71]. Further, climate and fungal symbionts of orchids may interact to shape the evolutionary response of specific vital rates to climate change, such as sprouting after dormancy [72].

### Implications for conservation

This study demonstrates the need for additional and more frequent surveys of rare plant populations to improve the reliability of information in databases used by land management agencies. Land managers who make decisions on how best to conserve rare species often base their decisions in part on the abundance and distribution of those organisms as reported in databases. However, many reported populations may no longer be extant. Managers should be aware that the number of populations of rare species in their databases is potentially an overestimate, and consider resurveying populations in databases to document their presence and condition, with priority given to older reports of small populations, especially those at low elevations or other areas with high climate vulnerability. Species like *C. fasciculatum* may be candidates for assisted migration [73-75] as their low-elevation populations experience extinction and if expansion or colonization at higher elevation locations does not occur naturally. We suggest that development of propagation and planting techniques [e.g., 76-78] to allow for intervention is warranted, and needs to consider the fungal dependency of this rare orchid [79].

## Acknowledgments

The authors gratefully acknowledge the contributions and cooperation by the Medford District Bureau of Land Management, especially Bryan Wender and Mark Mousseaux, who compiled database records for the species regionally. Support was also provided the Rogue-Siskiyou National Forest and facilitated by Wayne Rolle and Barb Mumblo. Staff of the Institute for Applied Ecology assisted with this project, including Carolyn Menke who made the maps used in this paper, and Amanda Stanley and Heather Root who provided statistical support.

